# Long-term survival and induction of operational tolerance to murine islet allografts through the co-transplantation of cyclosporine A eluting microparticles

**DOI:** 10.1101/2023.02.14.528345

**Authors:** Purushothaman Kuppan, Jordan Wong, Sandra Kelly, Jiaxin Lin, Jessica Worton, Chelsea Castro, Joy Paramor, Karen Seeberger, Colin C. Anderson, Gregory S. Korbutt, Andrew R. Pepper

## Abstract

One strategy to prevent islet rejection, is to create a favorable immune-protective local environment at the transplant site. Herein, we utilize localized cyclosporine A (CsA) delivery to islet grafts via poly(lactic-co-glycolic acid) (PLGA) microparticles to attenuate allograft rejection. CsA microparticles alone significantly delayed islet allograft rejection compared to islets alone (p<0.05). Over 50% (6/11) of recipients receiving CsA microparticles and short-term cytotoxic T lymphocyte-associated antigen 4-Ig (CTLA4-Ig) therapy displayed prolonged allograft survival for 214 days, compared to 25% (2/8) receiving CTLA4-Ig alone (p>0.05). CsA microparticles + CTLA4-Ig islet allografts exhibited reduced T-cell (CD4^+^ and CD8^+^ cells) and macrophage (CD68^+^ cells) infiltration compared to islets alone. We observed reduced mRNA expression of proinflammatory cytokines (IL-6, IL-10, INF-γ & TNF-α; p<0.05) and chemokines (CCL2, CCL5, CCL22, and CXCL10; p<0.05) in CsA microparticles + CTLA4-Ig allografts compared to islets alone. Long-term islet allografts contained insulin^+^ and intra-graft FoxP3^+^ T regulatory cells. Rapid rejection of third-party skin grafts (C3H) in islet allograft recipients suggested that CsA microparticles + CTLA4-Ig therapy induced donor specific operational tolerance. This study demonstrates that localized CsA drug delivery plus short-course systemic immunosuppression promotes an immune protective transplant niche for allogeneic islets.

**Article Highlights:**

- Systemic immunosuppression limits patient inclusion for beta cell replacement therapies
- Localized islet graft immunosuppression may reduce drug toxicity and improve graft survival
- Cyclosporine eluting microparticles + CTLA4-Ig therapy induced donor specific operational tolerance
- Graft localized drug delivery can create an immune protective transplant niche

## Introduction

Islet transplantation is a proven strategy to restore glycemic control, reduce hypoglycemic unawareness, and stabilize HbA1c levels for a subset of patients with T1D (1). A long-term follow-up study has shown that graft survival, based on C-peptide concentrations (< 0.1 nmol/L), was 48% at 20 years after the first transplantation (2). Even though marked improvements have been made in clinical islet transplantation the reliance on chronic immunosuppression to protect islets from host auto and alloreactivity remains a significant obstacle to patient inclusion. These obligatory and often diabetogenic drugs pose inherit potential risks, including cytotoxic effects on vital organs, severe infections and tumorigenesis (3;4). To reduce systemic side effects, targeted strategies such as islet graft localized delivery of immunosuppressive agents is an attractive strategy to abrogate drug toxicity, improve β-cell survival and a possible adjuvant to achieving operational tolerance.

Herein, poly(lactic-co-glycolic acid) (PLGA) microparticles were fabricated for localized release of cyclosporine A (CsA) to allogeneic murine islet grafts. PLGA is a Food and Drug Administration authorized biodegradable and biocompatible polymer approved for numerous biomedical applications, including drug delivery (5). Our group has recently shown that co-localization of dexamethasone (Dex) eluting PLGA microparticles to islet allografts with CTLA4-Ig therapy yielded 80% of the recipients euglycemic for 60 days posttransplant compared to 40% in empty microparticles + CTLA4-Ig treated recipients (6). CsA, a calcineurin inhibitor, is a potent immunosuppressive drug used to prevent allograft rejection in solid organ transplantation and to treat autoimmune diseases (7). In islet transplantation, CsA has been administered systemically along with immune cell-depleting agents (Anti-Thymocyte Globulin (ATG), Alemtuzumab) to prevent islet graft rejection as it blocks interleukin (IL)-2–dependent proliferation and differentiation of T cells (8). Additionally, CsA binds with cyclophilin D, which prevents mitochondrial permeability transition pore opening under stress conditions, stabilizing mitochondrial functions and preventing cell death (9). Conversely, it is well documented that high doses of CsA impair β-cell function (10-14). In addition, its nephrotoxicity is a serious side effect that limits CsA’s widespread clinical application (15). Although delivery of CsA using PLGA micro-particles has been previously utilized (16; 17), the targeted and localized effect of CsA on islet transplantation remains unexplored.

Therefore, we sought to investigate the ability of localized CsA delivery via PLGA microparticle elution to improve islet allograft outcomes. Here, we described the optimization of CsA eluting PLGA microparticle fabrication and *in vitro* characterization. Next, we assessed the safety, and toxicity of CsA eluting PLGA microparticles using a syngeneic transplant model. Subsequently, we examined the ability of CsA microparticles to enhance murine islet allograft function in a fully mismatched model (minor and major histocompatibility complex; MHC). Finally, we examined the ability of another clinically relevant immunosuppressant, CTLA4-Ig, to work in concert with CsA microparticle co-delivery to improve alloislet engraftment and survival.

## Research design and methods

### Preparation and characterization of CSA eluting PLGA microparticles

The procedure for the preparation and characterization of CSA eluting PLGA microparticles can be found in Supplementary Material.

### *In vivo* drug release characterization

*In vivo* CsA release study was conducted in nondiabetic B6.129S7-Rag1^tm1Mom^/J mice (Jackson Laboratory, Canada) following implantation of 4 mg of CsA microparticles mixed with collagen type I (Corning, NY, USA) and implanted under the kidney capsule (KC). At 0, 1, 3, 7, 14 and 21 days post-implantation (n=4 mice/ time pt), microparticle-bearing kidneys were removed, homogenized, and analyzed for CsA using HPLC (Agilent Technologies, 1200 series, CA, USA)(6; 18).

### Islet isolation, transplantation and metabolic follow-up

Procedures for islet isolation, transplantation and metabolic follow-up can be found in Supplementary Material. Syngeneic islet transplantation studies (BALB/c to BALB/c) were conducted to assess the safety and toxicity of CsA microparticles, subsequently, allogeneic islet transplants (BALB/c to C57BL/6) were conducted. Transplants consisted of 500 islets combined with 4 mg CsA microparticles under the KC of diabetic mice. Details of all experimental groups can be found in Supplementary Material.

### Intra-islet graft proinflammatory cytokine analysis and gene expression analysis

Acute allotransplant studies were conducted where diabetic C57BL/6 mice received 500 BALB/c islets + 4 mg CsA microparticles (n=3) or 500 BALB/c islets + 4 mg CsA microparticles + CTLA4-Ig (10 mg/kg, day 0, 2, 4, and 6 posttransplant). After 7 days, grafts were removed and analyzed for proinflammatory cytokines and gene expression.Procedures for intra-islet graft proinflammatory cytokine analysis and gene expression analysis and details of all experimental groups can be found in Supplementary Material.

### Islet graft immunohistochemical analysis

At different time points post-transplant, islet grafts were fixed in formalin, processed and sections were immuno-histochemically stained. Histological procedures can be found in Supplementary Material.

### Assessment of tolerance induction by allogeneic skin graft transplantation

The procedure for the assessment of tolerance induction by allogeneic skin graft transplantation can be found in Supplementary Material. Briefly, skin grafts were conducted on long-term functioning islet allograft recipients, to investigate the possibly of the induction of transplant tolerance (19; 20).

### Statistical analysis

All data are presented as mean±standard error mean (SEM). Statistical significance between the treatment groups was calculated by one-way ANOVA (analysis of variances) and unpaired t-test. A Tukey posthoc test was used to analyze variances for multiple comparisons between the study groups. Kaplan-Meyer survival function of alloislet transplants and skin transplants were compared using the log-rank (Mantel-Cox) statistical analysis. A 95% confidence interval was used as a threshold for significant, p<0.05.

## Results

### CsA-microparticle preparation and characterization

CsA eluting PLGA microparticles were synthesized by a single emulsion solvent evaporation technique (Fig. 1). Scanning electron micrographs (SEM) revealed that CsA-loaded microparticles were smooth and spherical in shape (Fig. 1). The size distribution of the CsA-loaded particles exhibited an average size of 16.3±2.3 μm (Fig. 1). The total amount of CsA contained in the 10 mg of lyophilized PLGA CsA microparticles was 890.56±14.61 μg and the encapsulation efficiency was 89.02±1.46 %. *In vitro* CsA-microparticle release kinetics demonstrated that 100% of CsA was completely released from the particles within 30 days (Fig. 1). An *in vivo* CsA release study demonstrated that CsA release was more accelerated *in vivo* than in *in vitro*. CsA concentration in KC decreased over time compared to the initial loading (day 0). There was approximately 57.97±3.58 % of CsA released by day 21 *in vivo* (Supplementary Fig. 1A-B) whereas 22.70±0.69 % of CsA was released by the same time *in vitro* (Supplementary Fig. 1C).

**Figure 1:**
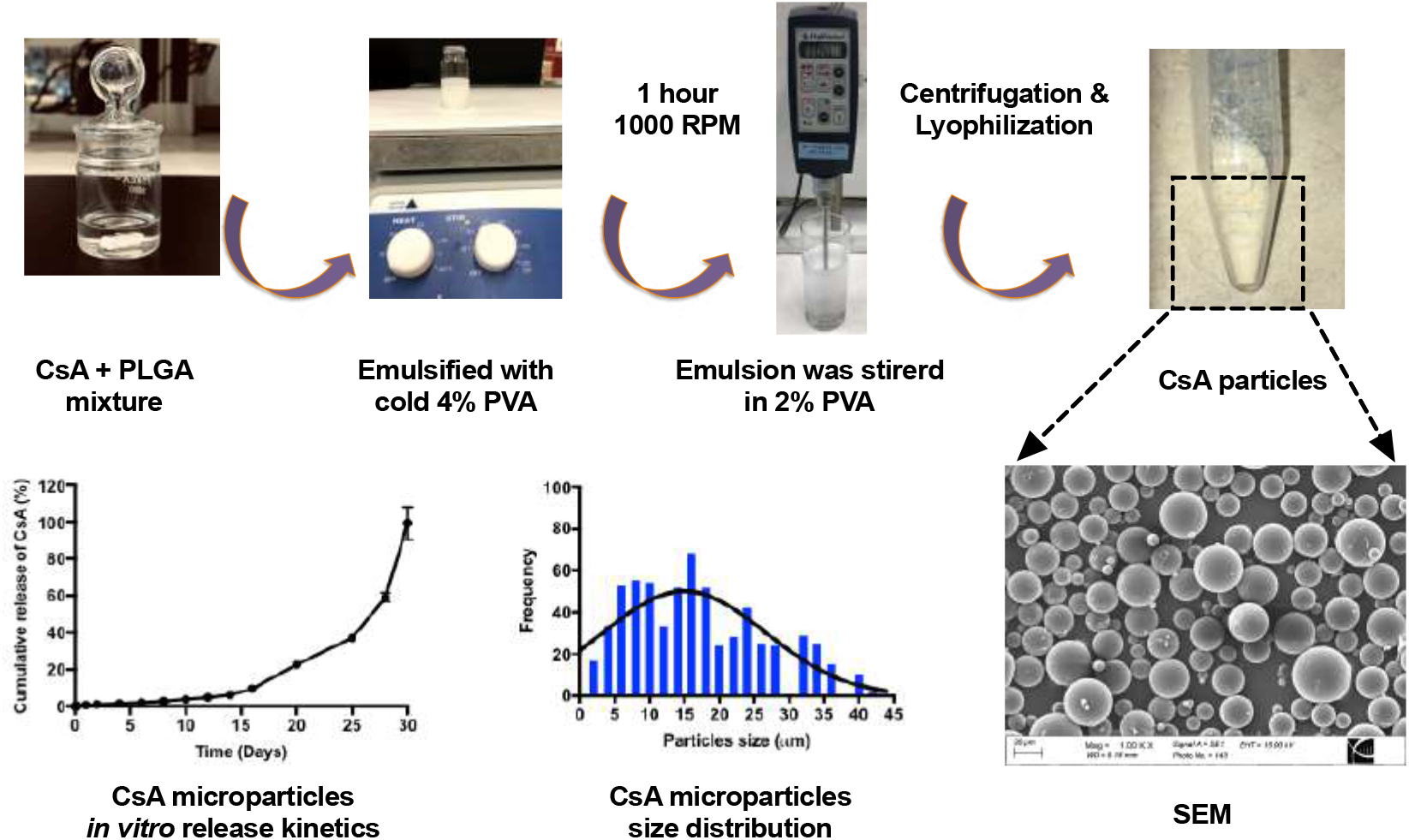
Schematic overview of the fabrication process for preparing cyclosporine A (CsA) eluting poly(lactide-co-glycolic acid) (PLGA) microparticles via a single emulsion (O/W) solvent evaporation technique. PLGA and CsA were dissolved in dichloromethane (DCM) followed by emulsified with cold polyvinyl alcohol (PVA). A scanning electron micrograph illustrates the surface morphology of CsA-loaded PLGA microparticles, and the histogram represents the size distribution of CsA particles. *In vitro* drug release characterization demonstrating the cumulative CsA release percentage (%) from PLGA-CsA particles over 30 days (n=4). Data points represent mean ± SEM. Scale bars represent 20 μm for the scanning electron micrographs.

### CsA eluting PLGA microparticles are non-toxic to murine syngeneic islet grafts

The adverse effects of localized CsA eluting microparticles were examined using a syngeneic mouse model, whereby BALB/c islets were co-transplanted with 4 mg of CsA microparticles (n=3) or islets alone (n=3) into the KC of diabetic BALB/c mice. SEM confirmed the co-localization of CsA-secreting microparticles on the surface of the islets (Fig. 2A). All mice co-transplanted with syngeneic islets alone (n=3, Fig. 2B) or syngeneic islets + CsA microparticles (n=3, Fig. 2C) became euglycemic within 2 days and maintained euglycemia throughout the follow-up period. At 35 days post-transplant, CsA microparticles + syngeneic islet recipients demonstrated a similar glucose clearance profile compared to islet alone recipients in response to a metabolic challenge (Fig. 2D and E); confirming that co-localization CsA microparticles did not impede islet engraftment or metabolic function, at this dose. The graft-bearing kidneys were removed 35 days post-transplant and subsequently, all recipients promptly returned to a hyperglycemic state, confirming graft-dependent euglycemia. Intact islets were observed in both CsA microparticles + islet grafts and islet alone grafts when stained for insulin & glucagon (Fig. 2F and H) and hematoxylin and eosin (H&E) (Fig. 2G and I). In addition, microparticles were observed as spherical vacuoles (indicated by *) within the CsA microparticles + islet grafts (Fig. 2I), but not in the islet alone grafts. These data demonstrate that 4 mg of lyophilized CsA microparticles (∼356 μg of CsA) are non-toxic and do not compromise islet graft function.

**Figure 2:**
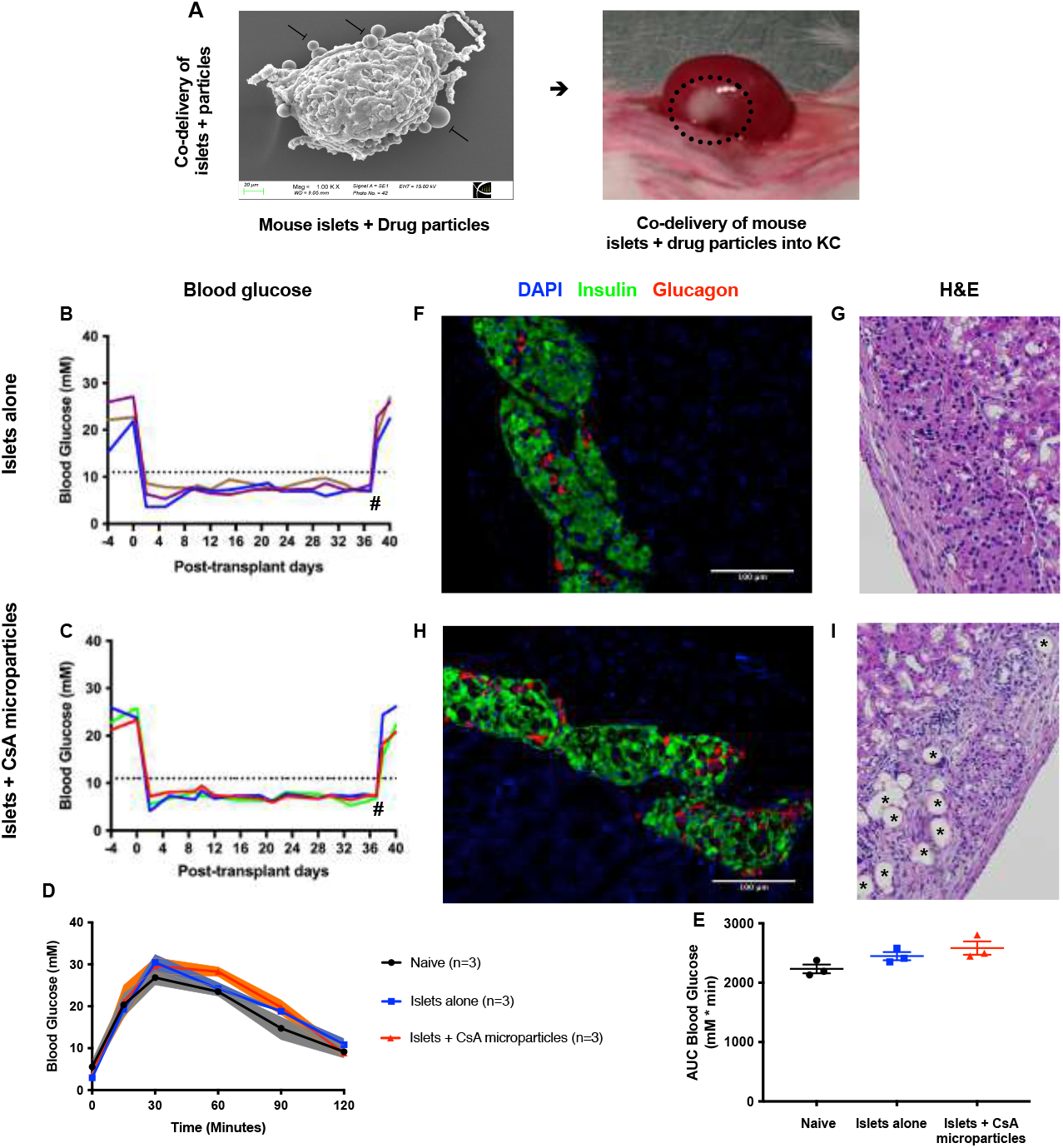
Syngeneic islet graft functional outcomes +/-CsA microparticle co-localization. When examined by SEM the microparticles were shown to be attached to the islet surface [A]. Diabetic BALB/c mice were transplanted with 500 BALB/c islets under the kidney capsule. Post-transplant blood glucose measurements of syngeneic islets alone (n=3) [B] and syngeneic islets + CsA microparticles (n=3) [C] recipients. Hash (#) represents the time of nephrectomy of graft bearing kidney. Intraperitoneal glucose tolerance tests of syngeneic islet alone recipients (n=3, blue) and syngeneic islets + CsA microparticles recipients (n=3, red), at 35 days post-transplant. Naïve nondiabetic, nontransplant mice served as controls (n=3, black). Mice were administered with 3 mg/g of 50% dextrose i.p. Blood glucose measurements were monitored at 0, 15, 30, 60, 90 and 120 min and analyzed for blood glucose [D] and area under the curve [E]. Immunohistochemistry of syngeneic islet alone grafts [F-G] and syngeneic islets + CsA microparticles grafts [H-I]. Green=insulin, red=glucagon and blue=nuclei. Asterisks (*) represented microparticles and were co-localized to islet grafts under the KC. Data were represented as mean ± SEM.

### Co-localization of CsA eluting microparticles prolongs murine allograft survival

We then conducted a series of fully allogeneic islet transplants [BALB/c islets (H2^d^) into diabetic C57BL/6 mice (H2^b^)]. Islet allografts containing CsA eluting microparticles (n=7) survived significantly longer than islet allografts co-transplanted with empty microparticles (n=8) (p <0.05) (Fig. 3A). Empty microparticles (control) recipients invariably lost graft function within 2 weeks posttransplant (median graft survival time (MST) = 9.87±1.66 days). In contrast, CsA microparticle recipients showed significantly improved graft survival (MST=19.00±2.64 days). To determine if CsA eluting microparticles could promote long-term islet allograft survival when combined with short-course systemic immunosuppression, we subsequently conducted a series of allografts ± CsA microparticles, whereby the recipients were administered with a low dose of CTLA4-Ig (10 mg/kg) i.p. at 0, 2, 4, and 6 days posttransplant. Recipients of CsA microparticles + CTLA4-Ig (n = 11) and empty microparticles + CTLA4-Ig (n = 8) displayed long-term allograft survival (Fig 3A). About 55% (6/11 recipients, MST = 147.70±23.81 days) of the recipients co-transplanted with CsA microparticles + islets + CTLA4-Ig maintained allograft function for 214 days post-transplant; doubling the long-term euglycemia prevalence compared to CTLA4-Ig alone monotherapy recipients (25%, 2/8 recipients, MST = 94.13±27.79 days). However, these data did not reach statistical significance (Fig. 3A, p>0.05). Both CsA microparticles + CTLA4-Ig treated alloislet recipients (n=7) and empty microparticles + CTLA-4-Ig (n=3) treated alloislet recipients demonstrated a similar glucose clearance profiles compared to naïve controls in response to a metabolic challenge (Fig. 3B-C); confirming that co-localization ± CsA microparticles did not impede islet engraftment or metabolic function, at this dose.

**Figure 3:**
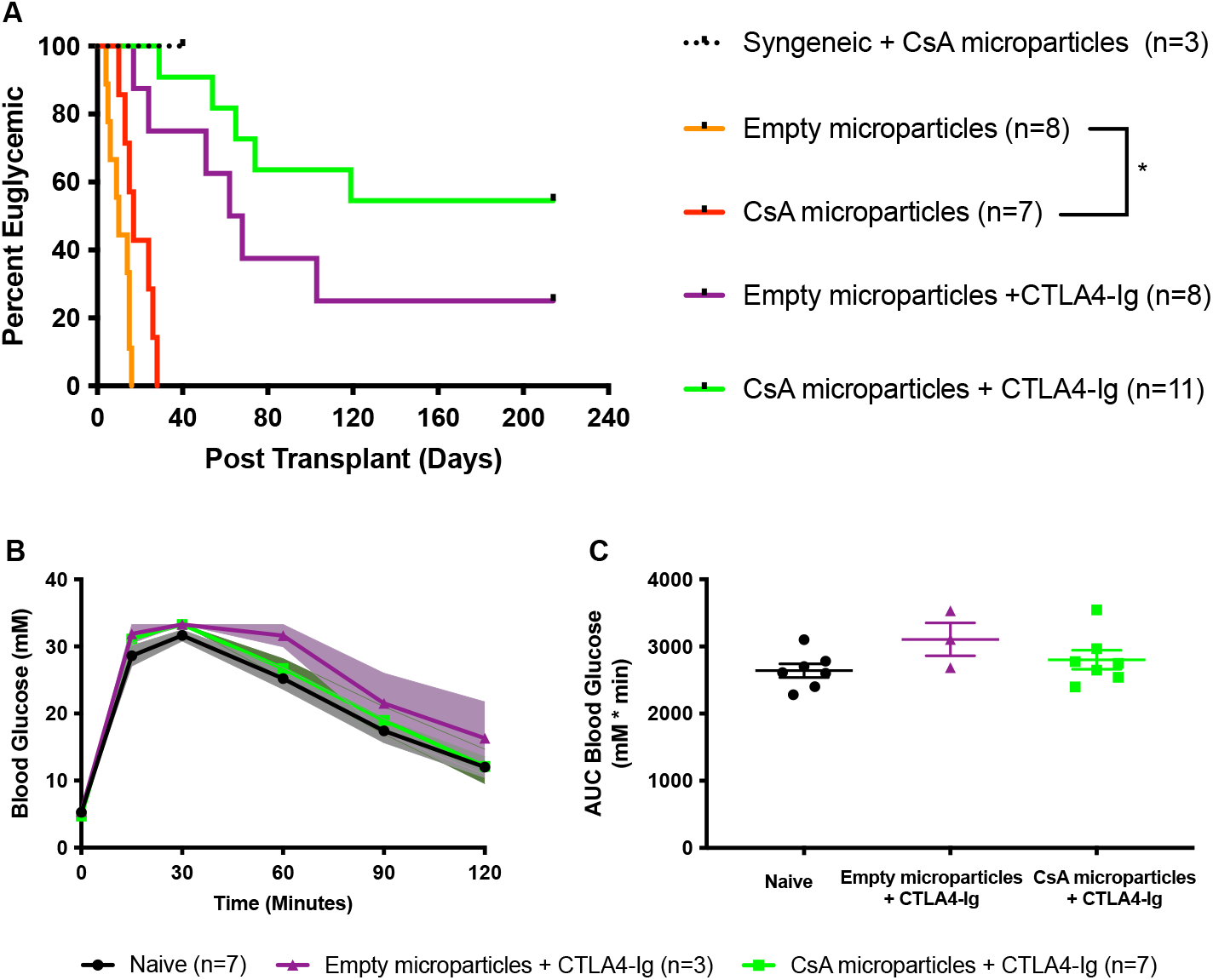
Islet allograft survival in mice transplanted with +/-CsA microparticles and +/-CTLA4-Ig [A]. Syngeneic islet graft + CsA microparticles (n=3, black) survival data were included as positive controls. Diabetic C57BL/6 mice were transplanted with 500 BALB/c islets under the KC. Islet allograft survival rates from recipients co-transplanted with islets + CsA eluting microparticles (n=7, red) significantly delayed the alloislet rejection compared to the islets + empty microparticles (n=8, orange) (p<0.05, Log-rank). Subsequently, two recipient groups (+/-CsA microparticles) were administered with CTLA4-Ig (10 mg/kg for short-term) i.p. at 0, 2, 4, and 6 days post-transplant. Recipients of empty microparticles + CTLA4-Ig (n=8, purple) demonstrated significant graft survival compared to empty microparticles alone (n=8, orange; p=0.0001, log-rank). Similarly, recipients of CsA microparticles + CTLA4-Ig (n=11, green) displayed significant graft survival compared to CsA microparticles alone recipients (n=7, red; p<0.0001, log-rank). The combination of CsA microparticles + CTLA4-Ig yielded a doubling in islet allograft survival up to 214 days compared to empty microparticles + CTLA4-Ig (55 vs 25%, respectively; p=0.13, log-rank). Allograft recipients that maintained euglycemia at 214 days post-transplant were electively subjected to survival nephrectomy to confirm graft-dependent function. Intraperitoneal glucose tolerance tests were performed on allogeneic euglycemic recipients of CsA microparticles + CTLA4-Ig (n=7, green), empty microparticles + CTLA4-Ig (n=3, purple) at 100 days post-transplant [B-C]. Naïve nondiabetic, non-transplanted mice served as controls (n=7, black). Mice were administered with 3 mg/g of 50% dextrose i.p. Blood glucose measurements were monitored at 0, 15, 30, 60, 90 and 120 min and analyzed for blood glucose profiles [B] and area under the curve [C]. Both CsA microparticles + CTLA4-Ig recipients (n=7, green) and empty microparticles + CTLA4-Ig recipients (n=3, purple) demonstrated a comparable glucose clearance to the naïve control (n=7, black; p>0.05, Anova). Data points were represented as mean ± SEM.

### Co-localization of CsA microparticles modulates intra-islet allograft proinflammatory responses and immune cell infiltration

To aid in elucidating the mechanisms by which CsA microparticles improve islet allograft survival, a series of acute allograft transplants were conducted and further analyzed by immunohistochemistry, proinflammatory cytokine secretion and inflammatory cell gene expression. Acute allogeneic islet grafts were stained for H&E, insulin & glucagon, CD4, CD8, CD68 and FoxP3. We observed qualitatively higher mononuclear cellular infiltration, into and surrounding the islets alone allografts (Fig. 4A1) compared to the CsA microparticles grafts (Fig. 4A2) or in the combination of CsA microparticles + CTLA4-Ig grafts (Fig. 4A3). Intact islets were observed in both CsA microparticles and CsA microparticles + CTLA4-Ig grafts; Fig. 4B2-B3). Qualitatively, both CsA microparticles and CsA microparticles + CTLA4-Ig grafts displayed a reduced presence of T cell populations such as CD4^+^ (Fig. 4C2-C3) and CD8^+^ (Fig. 4D2-D3) cells and macrophages (CD68^+^ cells) (Fig. 4E2-E3) compared to the islet alone grafts (Fig. 4C1, D1 and E1). Furthermore, intra-graft FoxP^3^ positive cells (T regulatory cells (Tregs)) were observed in the CsA microparticles and CsA microparticles + CTLA4-Ig treated grafts (Fig. 4F2-F3) but not in the islet alone grafts (Fig. 4F1). These results demonstrate that graft localized CsA release reduces the infiltration of mononuclear cells.

**Figure 4:**
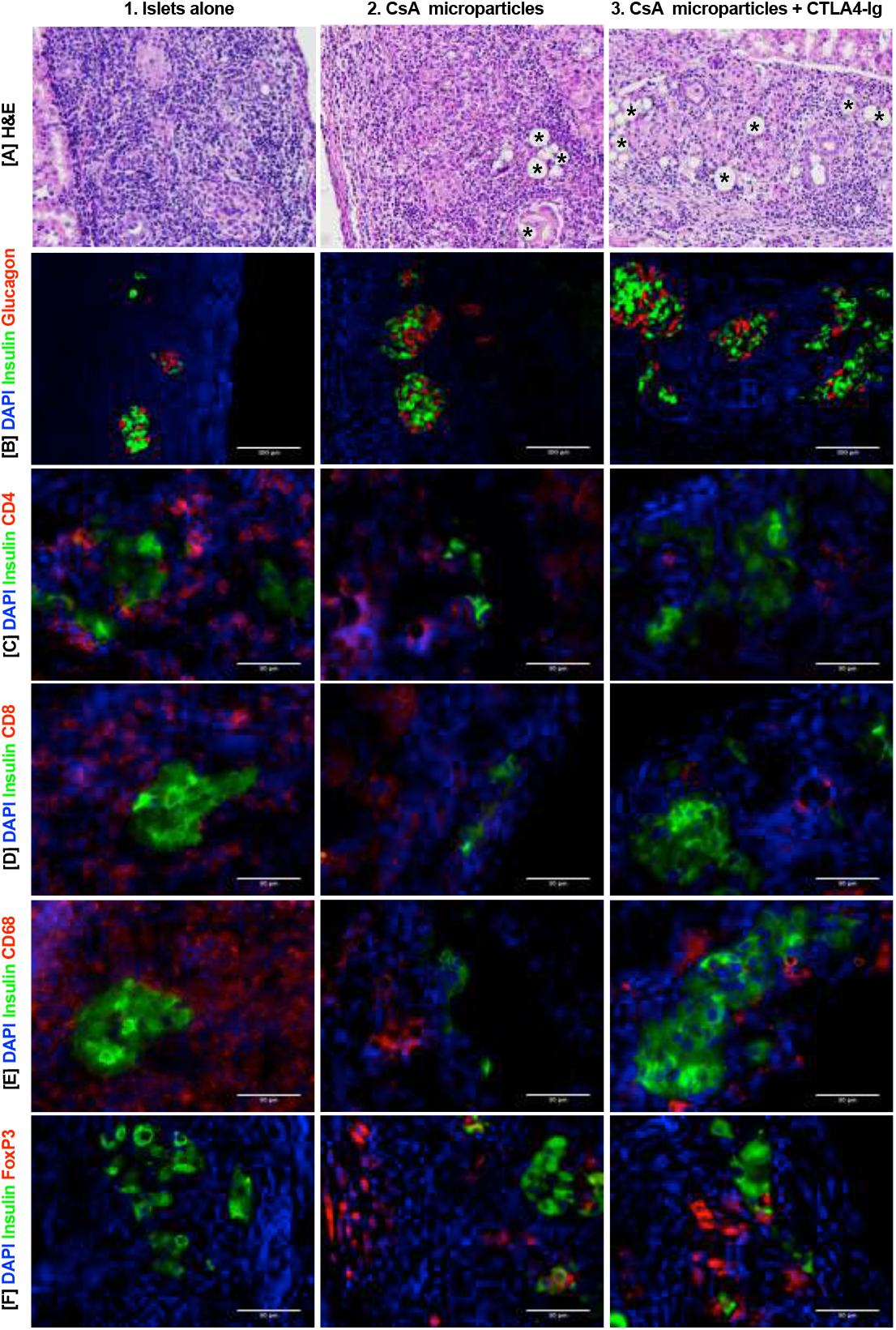
Immunohistochemistry of acute mouse islet allografts explanted 7 days post-transplant under the KC. Representative H&E staining of acute allogenic grafts revealed the presence of intact islet clusters with qualitatively minimal infiltration of mononuclear cells observed in CsA microparticles [A2] and CsA microparticles + CTLA4-Ig [A3] grafts, compared to islets alone grafts [A1]. Immunostaining of allogeneic grafts demonstrated that the intact islet clusters were positive for insulin (green) and glucagon (red) [B1-B3]. CsA microparticles and CsA microparticles + CTLA4-Ig treated grafts exhibited qualitatively minimal infiltration of CD4^+^ [C2-C3], CD8^+^ [D2-D3] and CD68^+^ [E2-E3] cells, while islets alone grafts showed higher infiltration of these mononuclear cells [C1, D1 E1]. FoxP3 (red) and insulin (green) immunostaining revealed the presence of intra-islet graft FoxP3^+^ (Tregs) cells in both CsA microparticles and CsA microparticles + CTLA4-Ig grafts [F2-F3], while islets alone grafts exhibited an absence of FoxP3^+^ cells [F1]. Scale bars represented as 50 μm and 100 μm. [Representative images, n = 2 per group and n=8 to 12 fields examined]

A cohort of acute allografts were analyzed for intra-graft proinflammatory cytokine profiles at 7 days posttransplant. IL-1β, IL-6, INF-γ and TNF-α expression trended towards being reduced in both CsA microparticles and CsA microparticles + CTLA4-Ig treated recipients compared to islet alone recipients (Fig. 5A-D). Of note, CsA microparticles + CTLA4-Ig grafts had significantly lower expression of TNF-α than islets alone grafts (p<0.05). IL-10, IL-12p70 and KCGRO expression was also measured; however, they were undetected in all grafts.

**Figure 5:**
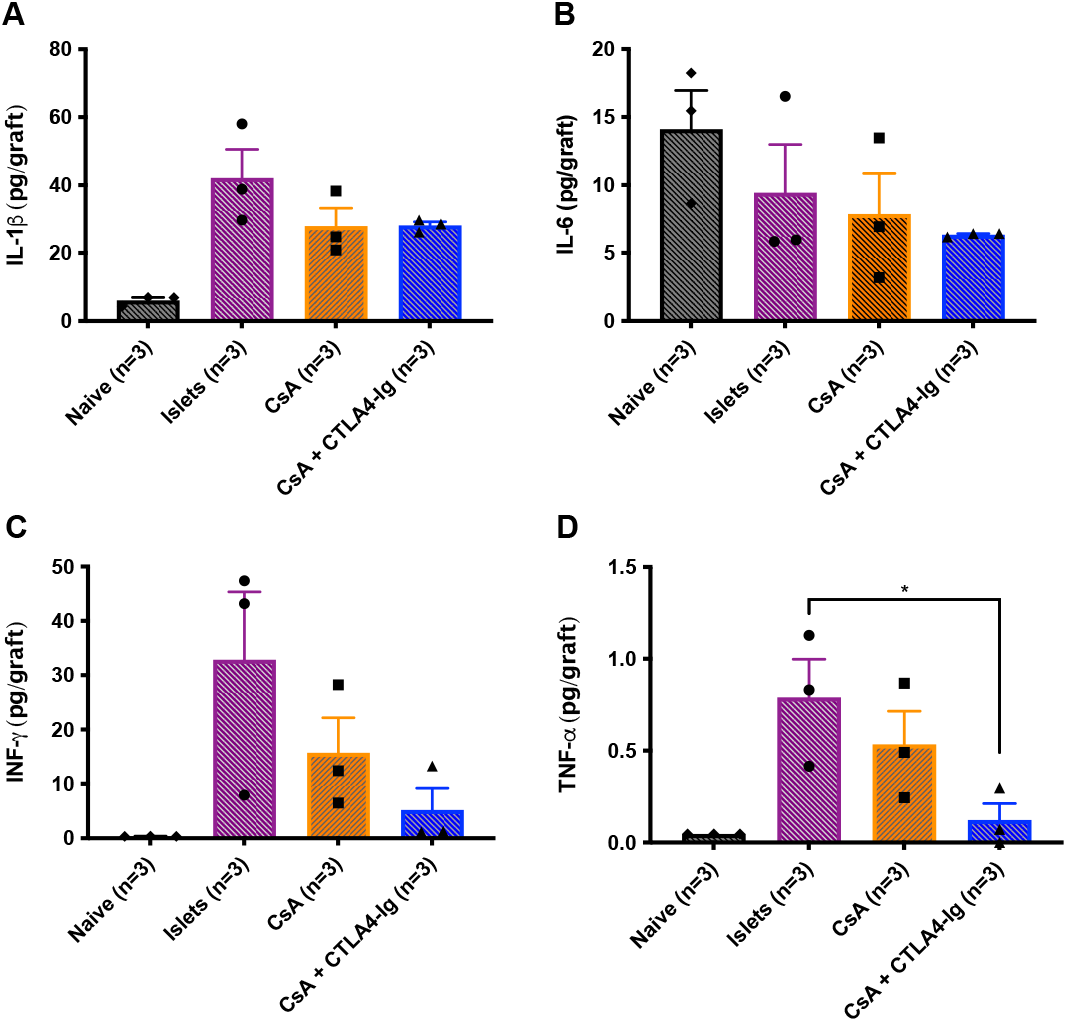
Intra islet graft pro-inflammatory cytokines analysis of acute allografts at 7 days post-transplant. Diabetic C57BL/6 mice were transplanted with BALB/c islets (500 islets) under the KC. Mouse recipient groups included were islets alone (n=3, purple), CsA microparticles alone (n=3, orange) and CsA microparticles + CTLA4-Ig (n=3, blue). Kidney from nondiabetic, non-transplanted C57BL/6 mice (n=3, black) served as a negative control. CTLA4-Ig (10 mg/kg) was administered i.p. at 0, 2, 4, and 6 days post-transplant. Grafts bearing kidneys were excised and homogenized to analyze the presence of pro-inflammatory cytokines. Proinflammatory cytokines such as IL-1β [A], IL-6 [B], INF-γ [C] and TNF-α [D] were observed at reduced concentration in CsA microparticles and CsA microparticles + CTLA4-Ig grafts compared to the islet alone grafts (p>0.05, Anova). CsA microparticles + CTLA4-Ig treated recipients showed significantly lower TNF-α concentration compared to islet alone recipients [D] (p<0.05, Anova). Cytokines like KC/GRO, IL-10 and IL-12p70 were not detected in the grafts (data not shown). Data points were represented as mean ± SEM.

A final cohort of recipients were analyzed for intra-graft mRNA expression of proinflammatory cytokines, chemokine and T-cell & macrophage markers using real-time RT-PCR at 7 days posttransplant. Reduced mRNA expression of proinflammatory cytokines (*IL-6, IL-10, INF-γ & TNF-α*) and proinflammatory chemokines (*CCL2, CCL5, CCL22, & CXCL10*) were observed in CsA microparticles treated recipients compared to the islet alone recipients (Fig. 6A-B). Most notably, *IL-6, IL-10, TNF-α, CCL2* and *CCL22* mRNA expression were significantly reduced in the CsA microparticles + CTLA4-Ig treated recipients compared to the islet alone recipients (Fig. 6A-B, p<0.05, p<0.01).

**Figure 6:**
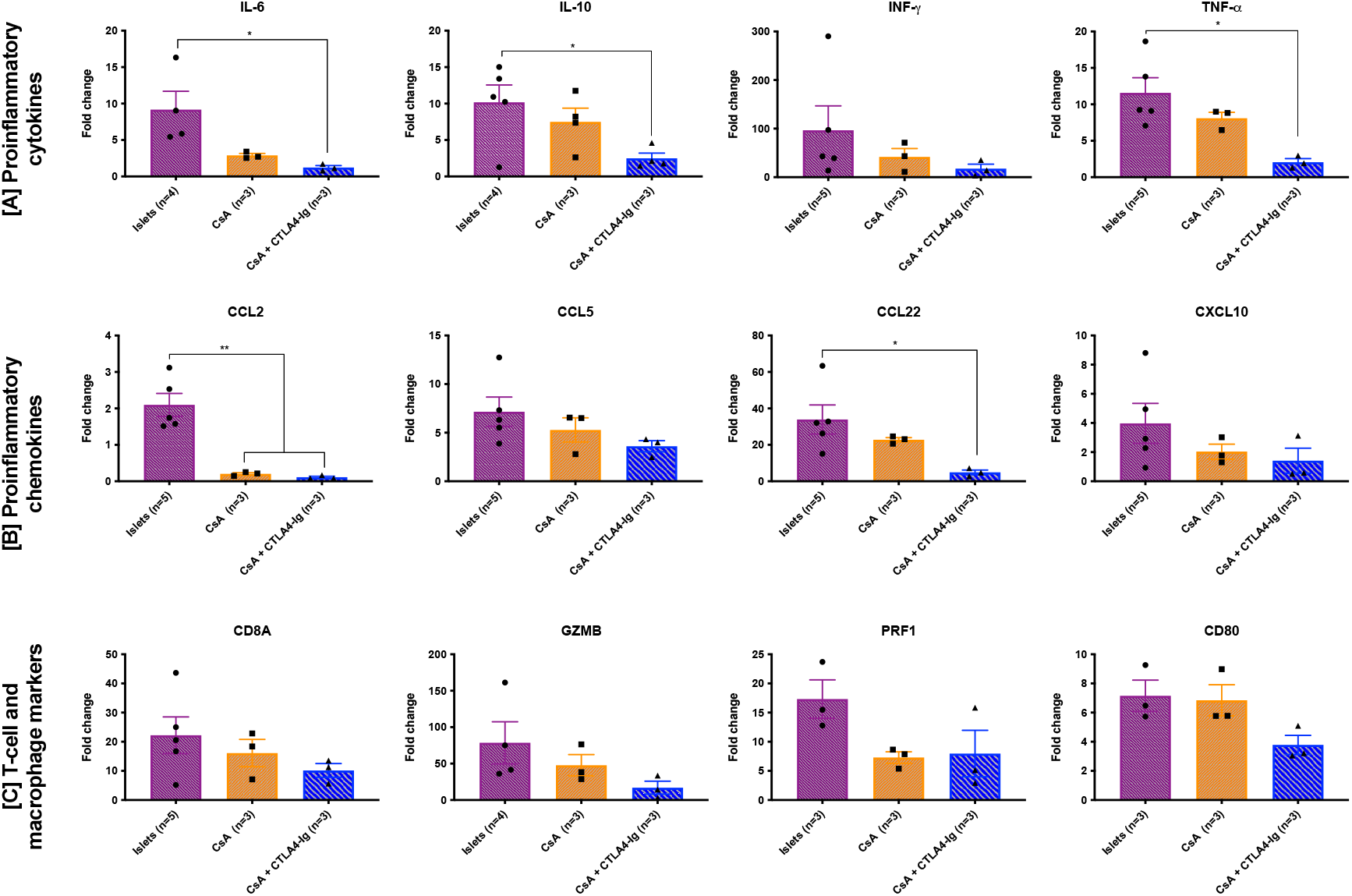
Gene expression from acute 7 days intra-islet grafts. Diabetic C57BL/6 mice were transplanted with BALB/c islets (500 islets) under the KC. Mouse recipient groups included were islets alone (purple), CsA microparticles alone (orange) and CsA microparticles + CTLA4-Ig (blue). Kidneys from sham transplanted C57BL/6 mice (n=3) served as a negative controls. CTLA4-Ig (10 mg/kg) was administered i.p. at 0, 2, 4, and 6 days post-transplant. CsA microparticles + CTLA4-Ig grafts showed significantly lower proinflammatory cytokine mRNA expression (IL-6, IL-10, and TNF-α except for INF-γ) compared to islet alone grafts [A] (p<0.05, Anova). Similarly, CsA microparticles + CTLA4-Ig treated grafts showed significantly lower proinflammatory chemokine mRNA expression (CCL2, and CCL22) compared to islet alone grafts [B] (p<0.05, p<0.01 Anova). However, CCL5 and CXCL10 mRNA expression was reduced in CsA microparticles + CTLA4-Ig grafts compared to islet-alone grafts [B] (p>0.05, Anova). T-cell markers (CD8A, GZMB, and PRF-1) and macrophage marker (CD80) mRNA expression trended towards being reduced in CsA microparticles + CTLA4-Ig grafts compared to islet alone grafts [C] (p>0.05, Anova). Data points were represented as mean ± SEM.

Furthermore, T cell markers (*CD8A, GZMB, & PRF1*) and macrophage marker (*CD80*) mRNA expression levels were modestly decreased in the CsA microparticles and CsA microparticles + CTLA4-Ig treated groups compared to the islet alone recipients (Fig. 6C, p>0.05).

### Histological characterization of long-term islets allograft

H&E staining of long-term grafts (214 days) from both CsA microparticles + CTLA4-Ig and empty microparticles + CTLA4-Ig demonstrated the presence of intact islets (Fig. 7A). Further, immunohistochemical staining demonstrated that the grafts were predominantly consisted of insulin-positive β-cells with few glucagon-positive α-cells (Fig. 7B). In addition, we further observed the presence of intragraft FoxP3 cells (Tregs) in both empty microparticles + CTLA4-Ig and CsA microparticles + CTLA4-Ig grafts (Fig. 7C-D).

**Figure 7:**
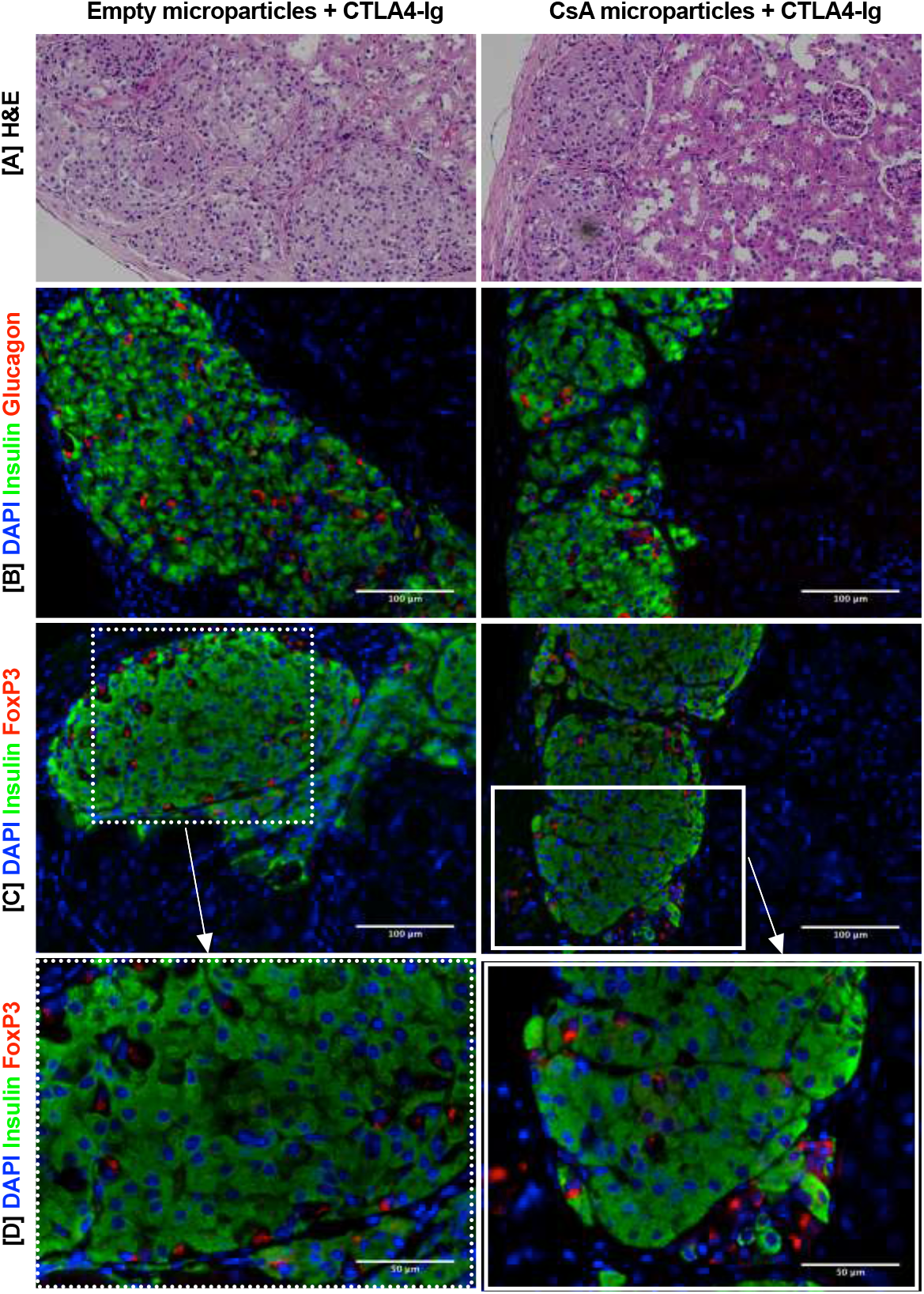
Immunohistochemistry of long-term islet allografts explanted electively at 214 days post-transplant under the KC. H&E staining revealed the presence of intact islet clusters in both CsA microparticles + CTLA4-Ig and empty microparticles + CTLA4-Ig grafts [A]. Immunostaining of long-term allogeneic islet grafts demonstrated that intact islet clusters were positive for insulin (green) and glucagon (red) in both CsA microparticles + CTLA4-Ig and empty microparticles + CTLA4-Ig grafts [B]. FoxP3 (red) and insulin (green) immunostaining revealed the presence of intra-islet graft FoxP3^+^ (Tregs) cells in both CsA microparticles + CTLA4-Ig and empty microparticles + CTLA4-Ig grafts [C-D]. Scale bars represented as 50 μm and 100 μm. [Representative images, n = 2 per group and n=10 to 13 fields examined]

### CsA microparticles + CTLA4-Ig treatment generated operational tolerance

To confirm that the immune tolerance achieved with CsA microparticles + CTLA4-Ig treatment is donor-specific, we performed skin grafts transplant on islet allograft recipients at 100 days posttransplant. We observed that BALB/c skin graft (donor-matched) rejection was significantly delayed in islet + CsA microparticles + CTLA4-Ig treated recipients (average skin graft rejection was 20.0±1.0 days) compared to the control recipients (non-islet transplanted and no CsA + CTLA4-Ig treated recipients, average skin graft rejection 9.57±0.20 days) (Fig. 8A, p<0.001, log-rank). In contrast, C3H skin (third-party skin) graft rejection was comparable between islet + CsA microparticles + CTLA4-Ig treated recipients (average skin graft rejection 11.83±0.74 days) and control recipients (non-islet transplanted and no CsA + CTLA4-Ig treated recipients, average skin graft rejection 10.29±0.28 days) (Fig. 8B). Syngeneic skin grafts were accepted [Supplementary Fig. 2A1]. We observed BALB/c skin grafts (donor-matched skin) [Supplementary Fig. 2B1] rejected gradually, whereas C3H skin (third-party skin) [Supplementary Fig. 2C1] rejected rapidly in recipients treated with islets + CsA microparticles + CTLA4-Ig. H&E staining of rejected BALB/c skin [Supplementary Fig. 2B2] and C3H skin [Supplementary Fig. 2C2] showed the absence of an intact epithelial layer compared to the naïve control skin [Supplementary Fig. 2B3 and C3] indicating the complete rejection of the skin grafts. Interestingly, alloislet transplanted recipients maintained normoglycemia until BALB/c skin graft rejection. The rejection of BALB/c skin appears to have triggered the rejection of the islet graft [Supplementary Fig. 3]. Rapid rejection of third party skin grafts together with delayed donor skin graft rejection indicated that graft localized CsA eluting microparticles + CTLA4-Ig promoted donor-specific immune tolerance.

**Figure 8:**
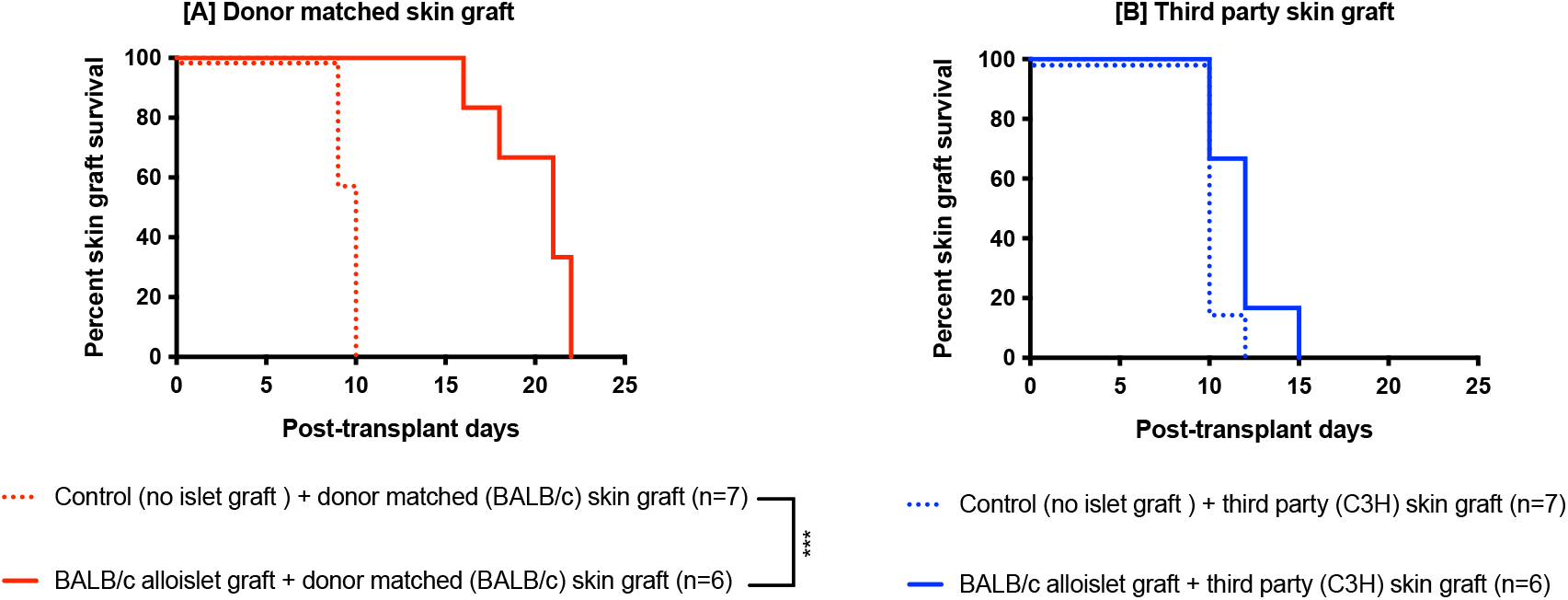
Skin graft transplants were conducted on islets + CsA + CTLA4-Ig treated recipients at 100 days post-transplant to investigate the transplant tolerance. Diabetic C57BL/6 mice were transplanted with BALB/c islets (500 islets) under the KC. CTLA4-Ig (10 mg/kg) was administered i.p. at 0, 2, 4, and 6 days post-transplant. Survival of skin grafts were analyzed on islets + CsA microparticles + CTLA4-Ig recipients, and control recipients (non-islet transplanted and no CsA + CTLA4-Ig treated recipients) [A-B]. BALB/c skin graft (donors matched skin) rejection was significantly delayed in islets + CsA microparticles + CTLA4-Ig treated recipients (average rejection 20.0±1.0 day) compared to the control recipients (non-islet transplanted and no CsA + CTLA4-Ig treated recipients, average rejection 9.57±0.20 day) (p<0.001, log-rank) [A]. In contrast, C3H skin (Third-party skin) graft rejection was comparable between islets + CsA microparticles + CTLA4-Ig treated recipients (average rejection 11.83±0.74 day) and control recipients (non-islet transplanted and no CsA + CTLA4-Ig treated recipients, average rejection 10.29±0.28 day)(p>0.05, log-rank)[B].

## Discussion

Allogeneic islet transplantation has successfully restored physiological glucose homeostasis in selected T1D patients (21). However, multiple factors such as oxidative stress, inflammatory insults, immune cells attack and systemic immunosuppression toxicity cause damage to the transplanted islets (22). In addition, the recurrence of islet autoimmunity may further accelerate the destruction of transplanted β-cells (23). Anti-inflammatory and immunosuppressive medications like etanercept, ATG, rapamycin, tacrolimus, and CsA are clinically used to prevent alloislet rejection. Nevertheless, these drugs are associated with undesirable side effects on the transplanted cells and the patient’s health (24; 25). Induction of local immunotolerance is an attractive strategy to prevent islet allograft rejection, which may improve graft function and broaden cellular therapy’s applicability while minimizing adverse side-effects of systemic immunosuppression (6; 26; 27). Several strategies have been established to evoke immunoprotection by co-delivering anti-rejection drugs with particles/scaffolds to the islet grafts (3; 6; 28-30). Immunomodulating drug-loaded microparticles provide multifaceted tools to locally modulate immune responses and represent an exciting tactic to aid cell transplantation applications. Furthermore, it is possible to load multiple drugs into a single carrier to target various pathways of the immune system simultaneously and adjust drug release kinetics by modifying the co-polymer ratios. Local release of therapeutic drugs is favoured over systemic administration, as a means to offset the harmful or adverse immune-dampening effects when left to circulate systemically (5; 23). Therefore, we developed and characterized the impact of islet graft localized CsA eluting PLGA microparticles on modulating the inflammatory responses in the islet transplant milieu.

CsA is a calcineurin inhibitor and its potent immunosuppressive property has profoundly improved solid organ transplant outcomes (31). CsA exerts immunomodulatory effects by blocking interleukin (IL)-2–dependent proliferation and differentiation of T cells and moreover, cyclosporine A - cyclophilin D complex formation stabilizes mitochondrial function and prevents cell death (8; 9). However, the diabetogenic effect and nephrotoxicity remain restricting the widespread use of CsA (17; 32). Furthermore, CsA exhibits low oral bioavailability owing to its poor biopharmaceutical properties such as its low aqueous solubility and low permeability which pose challenges in making a safe and effective delivery system (33). Consequently, there is a need for the development of a novel formulation strategy to encapsulate CsA with better bioavailability and fewer side effects. To the best of our knowledge, our study is the first to show the effect of CsA encapsulated PLGA particles in the context of islet transplantation. Drug encapsulation efficiency is affected by many factors including processing conditions, surfactants, solvents, molecular weight, surface functionalization, loading concentrations, and the nature of the drug and its interaction with the PLGA (34). We previously observed poor Dex encapsulation efficiency in PLGA microparticles (14.6 ± 1.7%) due to poor interactions between the Dex and PLGA (6). However, in this current study, we achieved improved CsA encapsulation efficiency of 89.02 ± 1.46 % in PLGA microparticles by modifying our emulsion process. We used a pre-cooled PVA solution to make the emulsion, this pre-cooling of PVA prevents solvent evaporation during the emulsion process. In addition, CsA is a cyclic peptide that has free functional groups had higher affinity interaction with the PLGA. Our observation validates earlier studies which demonstrated that the interaction of drug and polymer has a pivotal role in achieving a better drug encapsulation (34; 35).

Our data confirmed that a single administration of 4 mg of CsA microparticles (∼356 μg of CsA) to the localized graft site is safe, did not comprise islet graft function and yielded allograft protection over 5 weeks as compared to the allografts containing empty microparticles (no CsA). These findings contrast to when 50 mg/kg body weight of CsA was administered daily over three weeks in rats, which resulted in impaired glucose tolerance, a decreased insulin content and a decreased β-cell volume (12). However, we did not observe a durable graft function with this monotherapy, mirroring the observations of Arita *et al*., who showed that neither pravastatin nor CsA alone treated recipients prolonged alloislet graft survival (32).

Therefore, we speculated that combining localized CsA microparticles with other immunomodulatory delivery modalities (ie: CTLA4-Ig) would potentiate allograft protection. CTLA4-Ig effectively attenuates T-cell activation by competing with the costimulatory molecule CD28 to bind with ligands CD80 and CD86 on antigen-presenting cells (APCs), for which it has a higher affinity and avidity than CD28 (36). Indeed, previous islet allograft studies demonstrated that CTLA4-Ig costimulatory blockade alone yielded prolonged graft survival in approximately one-third of the murine recipients (37; 38). In a large animal non-human primate study, CTLA4-Ig therapy was demonstrated to prolong islet allograft survival in 40% of the recipients while demonstrating CTLA4-Ig’s effectiveness in suppressing both humoral and cellular immune responses (39). Islet allograft survival and euglycemic percent rate can be augmented when CTLA4-Ig is combined with other immunosuppressive agents such as Basiliximab (anti-IL-2R mAb), Sirolimus, and 3A8 (anti-CD40 mAb) (40).

Recently, our group demonstrated that co-localization of Dex microparticles + islet graft resulted in prolonged allograft function, when combined with low dose (10 mg/kg) administration of CTLA4-Ig i.p. (6). Therefore, we examined the ability of CTLA4-Ig to work in concert with CsA microparticle co-delivery to improve allograft survival. Mice that received allogeneic islets + CsA microparticles co-localization and short-term treatment of CTLA4-Ig showed 54% of the recipients maintained normoglycemia up to 214 days. In contrast, empty microparticles + CTLA4-Ig treated recipients exhibited only 25% of the recipients maintained normoglycemia for up to 214 days. CsA microparticles plus CTLA4-Ig treated recipients had doubled the long-term euglycemia prevalence compared to empty microparticles + CTLA4-Ig monotherapy recipients.

We hypothesized that localized CsA delivery to the islet grafts site increases primarily ß-cell survival by favourably modulating the transplant inflammatory milieu. It is well known that inflammatory mediators such as pro-inflammatory cytokines, pro-inflammatory chemokines, and complement activation products contribute to the early loss of islets and are negatively impacted islet graft functions (41). We observed that T-cells (CD4^+^ and CD8^+^ cells) and macrophages (CD68^+^ cells) populations were qualitatively reduced in our CsA microparticles or CsA microparticles + CTLA4-Ig treated recipients than the islet alone recipients. Previous studies demonstrated that the migration of activated CD8^+^ T cells to the islet grafts site triggers more damage and eventually destroys the transplanted allogeneic islet by releasing granzyme B (GZMB) and perforins (PRF1) (42). While CD4^+^ T cells indirectly contribute to graft rejection by boosting CD8^+^ T cells and secreting more inflammatory cytokines, such as interferon-gamma (IFN-γ) and tumour necrosis factor-alpha (TNF-α) (26). Our study also concurs with earlier observations that CsA microparticles + CTLA4-Ig treated recipients showed a marked reduction of mRNA expression of intra-islet graft cytokines (*IL-1ß, IL-6, IL-10, INF-γ and TNF-α*) and proinflammatory chemokines (*CCL2, CCL5, CCL22 and CXCL10*) known inducers of ß-cells apoptosis (43; 44). Similarly, proinflammatory chemokines (CXCL10, CCL2, CCL5 and CCL22) elicit poor islet allograft function (45; 46). These chemokines play an essential role in T1D disease progression and the islet graft rejection (27). We also observed reduced expression of *CD8A, GZMB, PRF1 and CD80* in CsA microparticles + CTLA4-Ig treated recipients. Thus, these observations prove that our CsA microparticles + CTLA4-Ig approach plays a vital role in altering the localized immune response while yielding improved allograft survival.

We observed long-term islet allograft function in CsA microparticles + CTLA4-Ig treated recipients. To determine if systemic tolerance was induced in these recipients, we performed skin grafts comparing to control recipients (non-transplanted and no CsA treated). We observed that BALB/c skin graft (donor-matched skin) rejection was significantly delayed in islets + CsA microparticles + CTLA4-Ig treated recipients compared to the control recipients. However, C3H skin (third-party skin) rejection was comparable between islets + CsA microparticles + CTLA4-Ig treated recipients and control recipients. Our skin grafts results are consistent with previous reports whereby naïve B6 mice rejected BALB/c skin allografts at 14 days posttransplant whereas intrahepatic islet transplanted tolerant B6 mice showed a modest prolongation of skin allograft survival for up to 17 days posttransplant (19). Our observations, align with others that reported that tolerance to Fas ligand chimeric with streptavidin (SA-FasL)-engineered islet grafts is donor-specific and systemic at the induction phase. Similar to our data, tolerance was antigen and tissue-specific, as SA-FasL-engineered islets failed to protect third-party islet and donor-matched skin (20). The rejection of donor-matched skin grafts, but not the third-party skin grafts, also culminated in the rejection of SA-FasL-engineered islet grafts. Similarly, in our study, we observed that alloislet recipients maintained normoglycemia until the BALB/c skin rejection indicating BALB/c skin graft rejection elicits vigorous allogeneic immune responses that simultaneously reject islet allografts (20). Taken together, our skin graft transplant findings suggest that our CsA microparticles + CTLA4-Ig treatments generated an operational tolerance.

We speculated that Tregs expression at the graft site could be causal for the observed transplant tolerance. It’s been reported that Treg cells traffic to allogeneic pancreatic islets immediately post transplantation in response to inflammatory cues, where they manifest their immunoregulatory function within the graft microenvironment (20). Moreover, Tregs play a crucial role in maintaining immune homeostasis and peripheral tolerance to foreign antigens in humans (47; 48). Also, it is reported that Tregs in the pancreas inhibits the actions of autoreactive T cells, thereby preventing diabetes progression (49). Our recent study also demonstrated that Dex eluting microparticles + CTLA4-Ig islet allograft contained more FoxP3^+^ Tregs. Indeed, in this present study, we observed the presence of intragraft FoxP3^+^ Tregs in both acute and long-term CsA microparticles + CTLA4-Ig islet allografts.

Collectively, our findings suggest that localized CsA drug delivery via microparticle elution provides a feasible therapeutic platform to deliver immunomodulatory drugs in a controlled and favourable manner at the islet graft site. This therapy is a safe and effective strategy for minimizing systemic immunosuppression and promotes durable islet allograft function with temporary systemic immunosuppressive therapy. Furthermore, CsA plus CTLA4-Ig treatment generated a donor-specific operational tolerance. We anticipate that this approach may be suitable for developing long-lasting immunomodulatory agents that effectively boost islet allograft survival without the need for systemic immunosuppression.

## Acknowledgements

We thank Lynette Elder from the Alberta Diabetes Institute Histology Core Lab, the University of Alberta for her technical assistance in the histochemical processing of the islet grafts.

## Author Contributions

PK contributed to the study design, preparation and characterization of drug-eluting microparticles, transplantation, data collection, analysis, and interpretation; and writing the manuscript. SK contributed to islet isolation and transplant studies. KS assisted with *in vitro* assays and PCR reactions. CC, JW, and JP helped with data collection. JW assisted with *in vitro* and *in vivo* release kinetics. JL and CCA contributed to skin graft studies. ARP and GSK contributed to the study design, data interpretation, critical editing of the manuscript and securing funding. GSK and ARP are the guarantors of this work, have full access to all the data in the study, and take responsibility for the integrity of the data and the accuracy of the data analysis.

## Prior Presentation

Parts of this study were presented in abstract form at the Virtual Congress of the International Pancreas and Islet Transplantation Association (IPITA), October 20-23, 2021 and Diabetes Canada and Canadian Society of Endocrinology and Metabolism (CSEM) Professional conference, Calgary, Canada, November 9-12, 2022.

