## Supplemental Files for "Long-term survival and induction of operational tolerance to murine islet allografts through the co-transplantation of cyclosporine A eluting microparticles"

### Supplementary Materials

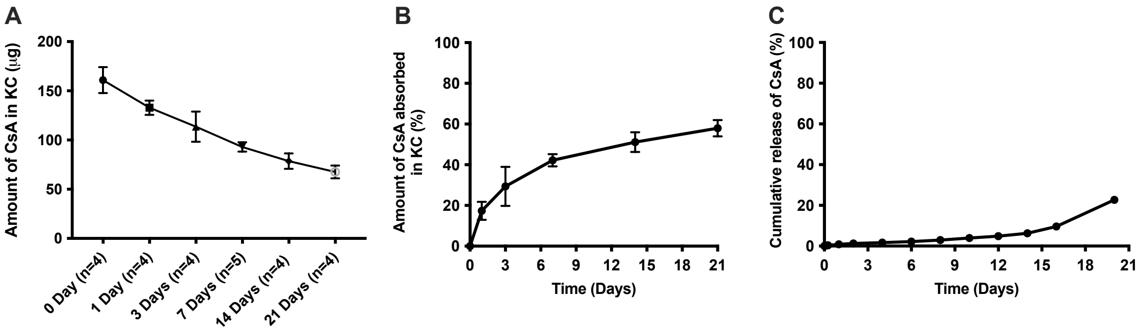


**Supplementary Figure 1:** *In vivo* release of CsA from CsA microparticles combined with collagen type I implanted under the KC over 21 days (A-B). *In vitro* release of CsA from CsA microparticles in PBS over 21 days (C).

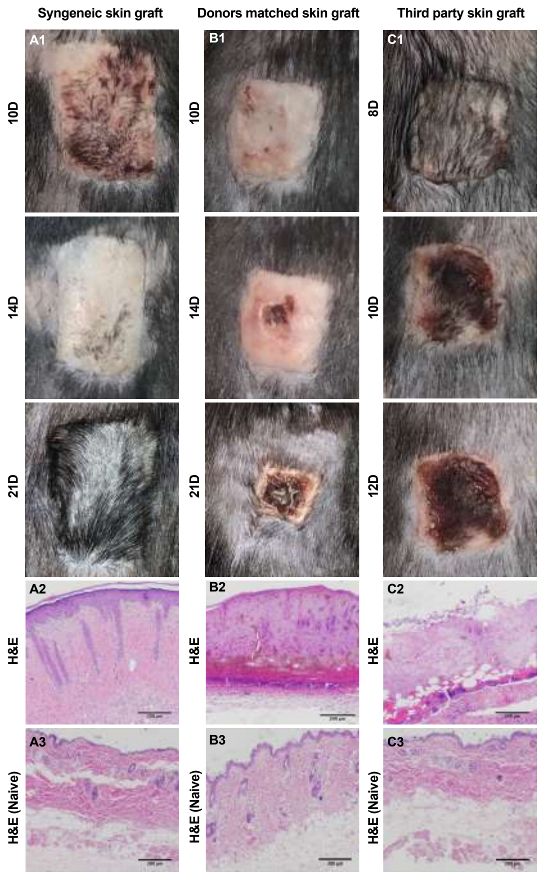


**Supplementary Figure 2:** Representative macroscopic images of skin grafts on mice treated with allogeneic BALB/c islets + CsA microparticles + CTLA4-Ig. Syngeneic skin graft (C57BL/6 mouse skin to C57BL/6 mouse) was accepted in recipient mice [A1], and also, observed an intact re-epithelialization in the accepted skin [H&E, A2]. Delayed BALB/c skin graft (donors matched skin) [B1] rejection was observed in recipients treated with islets + CsA microparticles + CTLA4-Ig compared to the C3H skin graft (Third-party skin) [C1] rejection in the same recipients. H&E staining of rejected BALB/c skin graft [B2] and C3H skin graft [C2] demonstrating the absence of an intact epithelial layer compared to the naïve control skin grafts [B3 and C3]. Scale bars; 200 μm.

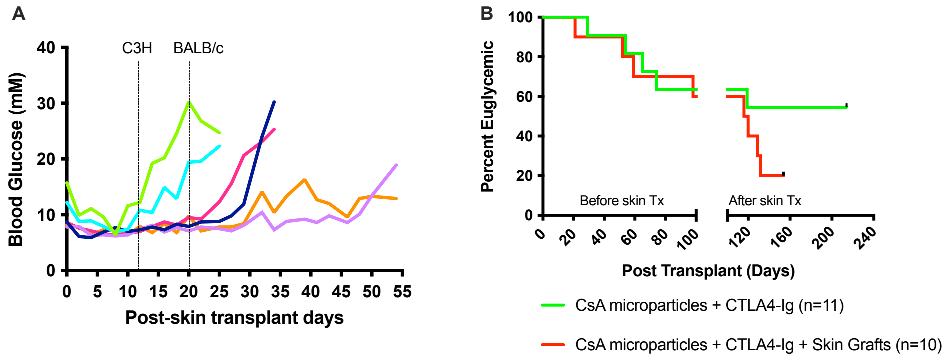


**Supplementary Figure 3:** Blood glucose levels of recipient mice treated with allogeneic BALB/c islets + CsA microparticles + CTLA4-Ig post-skin grafting. Black dotted lines demonstrate an average day of C3H (11.83±0.74 day) and BALB/c (20.0±1.0 day) skin graft rejection. Islet allograft survival, percentage euglycemia in mouse recipients transplanted with CsA microparticles + CTLA4-Ig before and after skin grafting [B].

`

**Supplementary Table 1** List of FAM-MGB (FAM: Fluorescein amidites, MGB: Minor groove binder) primer-probe mixes (Applied Biosystems-Thermo Fisher Scientific, MA, USA) was used in Real-time RT-PCR.

| **Gene Symbol** | **Assay ID** |
|---|---|
| IL-6 | Mm00446190_m1 |
| IL-10 | Mm01288386_m1 |
| INF-γ | Mm01168134_m1 |
| TNF-α | Mm00443258_m1 |
| CCL2 | Mm00441242_m1 |
| CCL5 | Mm01302428_m1 |
| CCL22 | Mm00436439_m1 |
| CXCL10 | Mm00445235_m1 |
| CD8A | Mm01182107_g1 |
| GZMB | Mm00442837_m1 |
| PRF1 | Mm00812512_m1 |
| CD80 | Mm00711660_m1 |

**Supplemental Materials**

**Preparation and characterization of CSA eluting PLGA microparticles**

Cyclosporine A (CsA, Mw 1202.6 g/mol, AdooQ Bioscience, CA, USA) eluting poly(lactic-co-glycolic acid) (PLGA: Mw 30,000-60,000 g/mol; 50:50 ratio, Sigma-Aldrich, ON, Canada) microparticles were prepared by a modified single emulsion (O/W, oil-in-water emulsion) solvent evaporation technique as previously described (1-3). Briefly, PLGA (5% weight/volume) and CsA (5 mg/mL) were dissolved in dichloromethane (DCM, Sigma-Aldrich, ON, Canada). In contrast to the Dexamethasone (Dex) eluting PLGA microparticles preparation, first, we used a single solvent system (Dichloromethane) to dissolve PLGA and CsA whereas a solvent mixture of dichloromethane (Sigma-Aldrich, ON, Canada) and acetone (Fisher Scientific, ON, Canada) at the ratio of 8:2 used to dissolve PLGA and Dex. Second, we increased PVA concentration from 0.5% to 2% and 4% to prepare CsA PLGA emulsion. Third, a new emulsion step introduced in the CsA PLGA emulsion process such as polymer-drug solution was emulsified using 10 mL of pre-cooled 4% polyvinyl alcohol (PVA, 80% hydrolyzed, Mw 9000-10000, Sigma-Aldrich, ON, Canada) solution with vigorous stirring for 5 minutes in a magnetic stirrer (VWR International, ON, Canada). Finally, all the contents were transferred into a 200 mL of 2% PVA solution and continuously stirred (high torque stirrer, Caframo®, BDC6015, ON, Canada) at 1000 rpm for an hour at room temperature. This process aids in completely evaporating the DCM while hardening the microparticle particles. Microparticles were collected by centrifugation (Allegra® X-15R centrifuge, Beckman coulter, NH, USA) at 3000 rpm for 5 minutes at 4°C followed by lyophilized (Dura-Dry™ MP, FTS Systems, NY, USA). Drug-free microparticles (empty microparticles) were prepared in the same manner as described above, however, CsA was not included in the solvent.

Empty PLGA microparticles were used in transplant studies to demonstrate their safety, tolerability and inability alone to confer protection from alloimmune responses.

CsA-loaded PLGA microparticle morphology was characterized by scanning electron microscopy (SEM, ZEISS EVO 10 Scanning Electron Microscope, Zeiss, NY, USA). Lyophilized CsA microparticles were spread onto carbon tape, coated with gold sputtering (Hummer 6.2 sputter coater, LADD Research Industries, VT, USA) and imaged. The particle diameter was measured manually using SEM images and a histogram was plotted. The amount of CsA entrapped within the microparticles (encapsulation efficiency) was quantified by high-performance liquid chromatography (HPLC) (Agilent Technologies, 1200 series, CA, USA) (4). Briefly, 10 mg of lyophilized CsA particles (n=15) were dissolved in acetonitrile (Sigma-Aldrich, ON, Canada): methanol (Fisher Scientific, ON, Canada) mixture (8:2) to release the CsA followed by centrifuged (Allegra® X-15R centrifuge, Beckman coulter, NH, USA) at 3000 rpm for 5 minutes. From this, a clear 10 μL solution was injected into the C 18 column, eluted at 0.25 mL/minute using acetonitrile: water (65:35) that detects the unknown CsA concentrations at 210 nm (4). The column temperature was maintained at 60°C. A series of known CsA concentrations between 10 μg/mL – 1000 μg/mL were run parallel to extrapolate the unknown sample concentrations.

*In vitro* drug release was conducted to understand the release pattern and kinetics of CsA microparticles (2; 3; 5; 6). In contrast to the Dex PLGA study, we did not use a dialysis bag to conduct the CsA release study and we also increased the microparticle concentration from 10 mg to 30 mg. Finally, we used a Multiskan SkyHigh microplate spectrophotometer to detect CsA concentration whereas Dex concentration was measured using ELSA. In brief, 30 mg of lyophilized CsA microparticles (n=4) were dispersed in 1 mL of phosphate-buffered saline (PBS, Thermo Fisher Scientific, ON, Canada) and maintained at 37°C. Two hundred microliters of samples were collected and replaced with the same amount of fresh PBS for 30 days. Samples were analyzed at 210 nm using a Multiskan SkyHigh microplate spectrophotometer (Thermo Fisher Scientific, ON, Canada) for the presence of CsA. A series of known CsA concentrations between 10 µg/mL – 100 µg/mL were run parallel to extrapolate the unknown sample concentrations.

**Islet isolation, transplantation and metabolic follow-up**

All animal studies were in accordance to the Canadian Council of Animal Care and guidelines obtained from the institutional ethical committee of the University of Alberta. All mice were maintained in a clean, sterile and pathogen-free environment and had access to *ad libitum* of water and pelleted food. All donors were male BALB/c mice (The Jackson Laboratory, Canada) between 8 and 12 weeks and weighed between 22 and 27 g. Mouse islets isolation and purification were conducted based on the previously described methodology (7). Briefly, collagenase (5 mg/mL) and thermolysin (0.2 mg/mL) (Liberase™ TL Research Grade, Roche Diagnostics, Mannheim, Germany) in Hank's Balanced Salt Solution (HBSS, Sigma-Aldrich, ON, Canada) was injected through the common bile duct to distend the pancreas *in situ* while blocking the duct entering at the duodenum. The distended pancreata were subsequently digested at 37°C for 14 minutes in a shaking water bath (Thermo Fisher Scientific, ON, Canada) at 50 rpm. Islets were purified from the pancreatic digest through histopaque density gradient (1.108, 1.083, and 1.069 g/mL, Sigma-Aldrich, ON, Canada) centrifugation at 3000 rpm for 12 minutes. Collected islets were washed with HBSS and cultured in Connaught Medical Research Laboratories (CMRL-1066, Mediatech,

Manasses, VA) supplemented with fetal bovine serum (10%), L-glutamine (100 mg/L), penicillin (112 kU/L), streptomycin (112 mg/L), and HEPES (25 mmol/L) at pH 7.4 for an hour prior to transplantation.

Prior to transplant, recipient male BALB/c and C57BL/6 mice (The Jackson Laboratory, Canada), between 8-12 weeks of age, were injected with 180 mg/Kg streptozotocin (STZ, Sigma-Aldrich, ON, Canada) reconstituted in acetate buffer (pH 4.5), intraperitoneally (i.p) to induce diabetes (3; 8). Animals with blood glucose readings >18 mmol/L, measured using OneTouch UltraMini glucose meter (LifeScan, Burnaby, BC, Canada), for two consecutive days were considered diabetic and utilized for subsequent transplant studies. Syngeneic islet transplantation studies were conducted to assess the safety and toxicity of CsA microparticles. Diabetic BALB/c mice were transplanted with either 4 mg (~ 356 μg of CsA) of CsA microparticles + 500 BALB/c islets (n=3) or islets alone (n=3) under the KC (7). A separate cohort of allogeneic mouse islets transplants was conducted by co-delivering 500 BALB/c islets with 4 mg CsA microparticles under the KC of diabetic C57BL/6 mice. Allogeneic experimental groups were (1) empty microparticles (no CsA) + islets (n = 8), (2) CsA microparticles + islets (n = 7), (3) empty microparticles + islets + CTLA4-Ig (n = 8), and (4) CsA microparticles + islets + CTLA4-Ig (n = 11). CTLA4-Ig (10 mg/kg, Biocell, West Lebanon, NH) was administered i.p on days 0, 2, 4, and 6 posttransplant. After transplantation, the recipient's blood glucose was monitored three times per week. Allograft rejection was defined as two consecutive blood glucose readings greater than 18.0 mM. In both syngeneic and allogeneic studies mice that maintained normoglycemia at 35 days and 100 days, posttransplant respectively were subjected to an intraperitoneal glucose tolerance test (IPGTT). Naïve, nondiabetic C57BL/6 (n = 7) and BALB/c (n = 3) mice served as controls. Overnight fasted mice were injected 3 g/kg of glucose (DMVet, Coaticook, QC, Canada) i.p. and blood glucose was measured at 0, 15, 30, 60, 90 and 120 minutes. Glucose clearance was expressed and compared by blood glucose area under the curve (AUC) values. Subsequently, allograft recipients that maintained euglycemia up to 214 days post-transplant were electively subjected to survival nephrectomize to confirm graft-dependent function. For histological assessment, graft-bearing kidneys were immediately fixed in 10% formalin (BDH Laboratory Supplies, VWR International, AB, Canada). Once recipients reverted to a pre-transplant hyperglycemic level, confirmed by two consecutive days, animals were euthanized.

**Intra-islet graft proinflammatory cytokine analysis and gene expression analysis**

Acute allotransplant studies were conducted where diabetic C57BL/6 mice received 500 BALB/c islets + 4 mg CsA microparticles (n=3) or 500 BALB/c islets + 4 mg CsA microparticles + CTLA4-Ig (10 mg/kg, 0, 2, 4, and 6 posttransplant). Diabetic mice that received islets alone served as a positive control (n=3) and non-transplanted diabetic C57BL/6 mice (n=3) served as a negative control. After seven days, graft-containing kidneys were removed and analyzed for intraislets graft proinflammatory cytokines via Mouse Pro-inflammatory 7-Plex Ultra-Sensitive Kit (Meso Scale Diagnostics, Rockville, MD) (3; 7).

Acute allotransplant studies were conducted where diabetic C57BL/6 mice received 500 BALB/c islets alone (n=3), or 500 BALB/c islets + 4 mg CsA microparticles (n=3) or 500 BALB/c islets + 4 mg CsA microparticles + CTLA4-Ig (10 mg/kg, 0, 2, 4, and 6 posttransplant). Naïve, sham transplanted (no islet graft) KC of C57BL/6 mice (n=3) was used to calculate the fold change of gene expression. After 7 days, the graft-containing kidneys were removed, processed with Trizol

(Thermo Fisher Scientific, ON, Canada) and cryopreserved at -80°C. Total RNA was extracted using the RNeasy Mini Kit (Qiagen, Mississauga, ON Canada) as per the manufacturer's protocol, followed and 1 μg of RNA was reverse transcribed into cDNA using the High Capacity cDNA Reverse Transcription Kit (Applied Biosystems- Thermo Fisher Scientific, MA, USA). All validated FAM MGB primer-probe mixes (FAM: Fluorescein amidites, MGB: minor groove binder, Applied Biosystems TaqMan Gene Expression Assay, MA, USA) were listed in Supplementary Table 1. Subsequently, cDNA was amplified using the StepOnePlus™ Real-Time PCR system with StepOne software v2.3 (Applied Biosystems, MA, USA), in MicroAmp Fast Optical 96 well plates (Applied Biosystems, MA, USA), using TaqMan Fast Advanced Master Mix (Applied Biosystems, MA, USA). Quantitative gene expression values were determined by the δ−δ CT method (CT: Cycle threshold) and normalized with the housekeeping gene, β-actin, and the calibrator naïve BALB/c mouse kidney control (9; 10). Gene expression data were expressed as relative fold change to the sham transplant.

**Islet graft immunohistochemical analysis**

Formalin-fixed islets grafts were processed, embedded in paraffin block and sectioned at a thickness of 5 μm. Subsequently, grafts were stained with hematoxylin and eosin (H&E). For immunohistochemical analysis, tissue slides were rehydrated, antigen retrieved and blocked with 20% normal goat serum (NGS) (Jackson ImmunoResearch, PA, USA) for 1 hour. After blocking, sections were incubated with the respective primary antibodies of anti-guinea pig α-insulin (1:5, Agilent CA, USA), mouse anti-glucagon (1:5000, Sigma-Aldrich, ON, Canada) for an hour or rabbit anti-FoxP3 (1:200, Novus Biologicals, CO, USA) at 4°C for overnight, followed by 3 series of washing with 1x PBS. Subsequently, sections were incubated with respective secondary antibodies of goat anti-guinea pig Alexa fluor 488, goat anti-mouse Alexa fluor 594 or goat anti-rabbit Alex flour 594 for 1 hour at room temperature (1:200, Thermo Fisher Scientific, ON, Canada). After incubation, tissue sections were washed with 1xPBS, 3 times, and subsequently coverslipped with 100 μL of DAPI anti-fade reagent (Thermo Fisher Scientific, ON, Canada) and dried at room temperature in the dark.

To assess immune cells infiltration in the grafts, procured grafts were preserved in cryomatrix (Epredia™ Cryomatrix™ embedding resin, Thermo Fisher Scientific, ON, Canada), immediately frozen at -20°C and tissue samples were cut in consecutive 5-μm sections on a cryotome (Leica CM3050 S Cryostat, Leica Biosystems Inc. ON, Canada), air dried, and fixed with cold acetone for 2 minutes. Subsequently, sections were air-dried, immersed in 1X PBS and blocked with 20% NGS (Jackson ImmunoResearch, PA, USA). Next, sections were incubated with respective primary antibodies of rat anti-mouse CD4, rat anti-mouse CD8 and rat anti-mouse CD68 (1/200, Bio-Rad, CA, USA) for 1 hour, followed by incubated for an hour with goat anti-rat Alex flour 594 (1:200, Thermo Fisher Scientific, ON, Canada). Slides were washed three times with 1xPBS between the primary and secondary antibody incubations. Subsequently, sections were counter-stained with insulin (1:5, Dako, CA, USA) and mounted with DAPI anti-fade reagent (Thermo Fisher Scientific, ON, Canada).

**Assessment of tolerance induction by allogeneic skin graft transplantation**

Skin grafts were conducted on long-term functioning islet allograft recipients, to investigate the possibly of transplant tolerance (11; 12). A separate cohort of alloislet transplant studies were conducted where diabetic C57BL/6 mice (n=10) received 500 BALB/c islets + 4 mg CsA microparticles + CTLA4-Ig (10 mg/kg, 0, 2, 4, and 6 posttransplant). Mice that maintained normoglycemia for 100 days received skin grafts from C57BL/6 (syngeneic grafts), BALB/c (islet donor-matched) and C3H (third-party). Naïve, non-islet transplanted and no CsA + CTLA4-Ig treated C57BL/6 mice received skin grafts similar to the experimental recipients. In brief, approximately 1 $cm^2$ of the donor's flank skin was transplanted at the left (BALB/c skin) and right (C3H skin) flank, and dorsally (between the flanks; C57BL/6 skin) and then secured with pressure bandages for 8 days. Subsequently, bandages were removed, and skin grafts were observed and scored every day to assess rejection. Skin grafts were considered rejected when less than 10% of viable tissue remained. The rejected and accepted skin grafts were collected and stained with H&E.
